# RNA-based therapy for *Cryptosporidium parvum* infection: proof-of-concept studies

**DOI:** 10.1101/2022.01.11.475978

**Authors:** A Castellanos-Gonzalez, A Sadiqova, J Ortega-Mendez, AC White

**Affiliations:** Department of Internal Medicine, Infectious Diseases Division, University of Texas Medical Branch, Galveston Texas, USA

**Author notes:** Corresponding author: Alejandro Castellanos-Gonzalez, Assistant Professor, Infectious Diseases Division, Internal Medicine department, University of Texas Medical Branch, Galveston Texas, USA.

**Keywords:** *Cryptosporidium*, Cryptosporidiosis, Gene silencing, Argonaute, NDK

## Abstract

*Cryptosporidium* is a leading cause of moderate-to-severe diarrhea in children. Nitazoxanide, the only FDA-approved treatment for cryptosporidiosis, has limited efficacy in those at highest risk for sequelae. RNA-argonaute (Ago) complexes to *Cryptosporidium* nucleoside diphosphate kinase (cpNDK) decreased the *Cryptosporidium parvum* mRNA by 95% in infected cells *in vitro*. Treatment of mice by oral gavage with ssRNA/Ago complexes encapsulated in lipid nanoparticles led to delivery of the complexes into intestinal epithelial cells. Treatment of *C. parvum* infected mice with ssRNA/Ago complexes targeting cpNDK led to the resolution of oocyst shedding in 4/5 SCID/beige mice. These results confirm the potential use of antisense therapy as an alternative approach to cryptosporidiosis treatment.

## Introduction

*Cryptosporidium* is a leading cause of moderate-to-severe diarrhea in children (1–3). Nitazoxanide is the only FDA-approved medication available for cryptosporidiosis treatment, but it has limited efficacy in malnourished children and is ineffective in immunocompromised individuals. More effective treatment options are urgently needed (4, 5). We have developed a method to silence genes in this parasite using pre-assembled complexes of *Cryptosporidium* single-stranded RNA and the human enzyme Argonaute 2 (ssRNA/Ago) (6, 7). We have used this method to determine the role of selected genes during *Cryptosporidium* infection (8–10).

Our studies indicated that silencing of Nucleoside diphosphate kinase (*NDK*), reduces proliferation and egress of *Cryptosporidium* parasites (8). NDK is a housekeeping enzyme that balances cellular nucleoside triphosphate (NTP) pools by catalyzing the reversible transfer of γ-phosphate from NTPs to nucleoside diphosphates (NDPs) (11). Microbial NDKs have roles in protein histidine phosphorylation, DNA cleavage/repair, and gene regulation (11). In Leishmania and microbial infections NDK has been linked with modulation of the host immune response (12, 13). Downregulation of NDK gene arrests cell proliferation in cancer cell lines (14). Previous studies have demonstrated the feasibility of delivering small interfering RNA (siRNA) to the mouse intestines and human epithelial cells to block expression of intestinal genes (15, 16). We hypothesized that ssRNA/Ago complexes may be used orally to treat *Cryptosporidium* infection. However, siRNA therapeutics face important drawbacks before use in the treatment of intestinal pathogens including siRNA stability in the intestine, off-target effects, successful siRNA delivery into the pathogen *in vivo*, and nonspecific activation of the host immune system (17). Here we evaluated stability of complexes during intestinal transit and delivery into infected cells. Also, we tested gene silencing on intracellular parasites and evaluated the anti-cryptosporidal effect of ssRNA/Ago complexes in a mouse model of cryptopsporidiosis.

## Methods

### ssRNA/Ago assembling and silencing experiments

We induced gene silencing in *Cryptosporidium* oocysts with ssRNA/Ago complexes or Naked ssRNA. For these experiments, we used capped 21-nt ssRNA (Integrated DNA Technologies, Cornville, AZ) and recombinant Argonaute (Ago2) protein as described (7). Briefly, ssRNA-hAgo2 complexes were assembled by combining 2.5 μl [100 nM] of ssRNA, 2 μl [62.5 ng/ul] of hAgo2 protein (Sino Biologicals, North Wales, PA), and 15 μl of assembling buffer (2 mM Mg (OAc)_2_, 150 mM KOHAc, 30 mM HEPES, 5 mM DTT, nuclease-free water). The mixture was incubated for 1 hr at room temperature. For parasite transfection, lyophilized protein transfection reagent [PTR] (Pro-Ject, Thermo Scientific, Rockford, IL) was reconstituted in 1 ml of HEPES 100 μm. The complexes or naked ssRNA were encapsulated by adding 15 μl of PTR suspension and incubating for 30 min at room temperature. To complete transfection, 5×10^5^ oocysts diluted in 5 μl of water were added to each reaction tube containing complexes in PTR and incubated at room temperature for 1 h. The slicer activity of hAgo2 was activated by incubating samples at 37 °C for 2 h. 21 nt NDK anti-sense or sense sequences were used to target NDK gene or as negative control (Supplementary Table 1).

### RNA extraction and evaluation of silencing by RT-PCR

Prior to RNA isolation, samples (previously stored at −20°C) were thawed at 95°C for 2 minutes. Total RNA was extracted using the Qiagen’s RNeasy Plus Mini Kit (Qiagen, Valencia CA). The RNA was eluted from purification columns with 100 μl of RNase-free water. The RNA concentration was determined by spectrophotometry using a NanoDrop 100 Spectrophotometer (Thermo Fisher Scientific, Waltham MA). Silencing in transfected oocysts was analyzed by qRT-PCR using qScript™ One-Step SYBR^®^ Green qRT-PCR Kit, Low ROX™ (Quanta BioSciences/VWR, Radnor, PA), using 2 μl of purified RNA template [20 ng/μl], 5 μl of the One-Step SYBR Green Master Mix, 0.25 μl of each primer at a 10 μM concentration, 0.25 μl of the qScript One-Step reverse transcriptase, and 4.25 μl of nuclease-free water for a total of 10 μl of mix per sample. The RT-PCR mixture (total volume 12 μl) was transferred to 96-well Reaction Plates (0.1 mL) and RT-PCR amplification conducted with a 7500 Fast Real-Time PCR System (Applied Biosystems, Foster City, CA) under the following cycling conditions: 50°C for 15 minutes, 95°C for 5 minutes, then 50 cycles of 95°C for 15 seconds and 63°C for 1 minute, followed by a melting point analysis (95°C for 15 seconds, 60°C for 1 minute, 95°C for 15 seconds and 60°C for 15 seconds). RT-PCR values were normalized to *Cryptosporidium* GAPDH (housekeeping gene). To calculate fold changes between NDK from control samples and NDK from silenced samples, we used the ∆∆Ct method (18). The primers used for RT-PCR are indicated in supplement table 2.

### Antisense treatment in vitro in HCT-8 cells and fluorescent microscopy

Human ileocecal cells (HCT-8 cells, ATCC, Manassas, VA) and *Cryptosporidium parvum* parasites (Waterborne, INC, New Orleans, LA) were used for in vitro studies. HCT-8 cells were cultured in 25 cm flasks with 5 ml of complete media (RPMI-1640 media [Gibco/Thermo Fisher Scientific, Waltham, MA] supplemented with 10% fetal bovine serum [Stemcell Technologies, Vancouver, Canada] and 1× antibiotic/antimycotic solution [Gibco/Thermo Fisher Scientific, Waltham, MA]) at 37 C. After overnight culture, HCT-8 cells were harvested and seeded (~5 × 10^5^ per well) onto cover slips placed in 6-well plates (Costar, Corning, NY) at 37 °C for 24 hrs. For excystation, oocysts were washed 3 times with 250 μL of phosphate-buffered saline (PBS), resuspended in 50 μL of acidic water (pH 2–3) and incubated for 10 minutes on ice. Excystation medium (complete medium supplemented with 0.8% taurocholate) was added to the sample, which was then incubated for 1 hour at 37°C. The released sporozoites were stained by adding 4 μl Carboxyfluorescein succinimidyl ester (CFSE). ~5 × 10^5^ Sporozoites were added to the HCT-8 cells (ratio 1:1). After 2 h, sporozoite media was removed, and 500 μl of complete media was added. The following day, labeled-ssRNA (ssRNA-Cy5 IDT) complexes were prepared and encapsulated in PTR. The labeled complexes in PTR were used to transfect infected cells for 24 hrs (at 37°C). After cell transfection, media was removed from chambers and cover slips were placed in petri dishes for washing (3 times rising with 200 μl of PBS). 1 ml of fixation solution (Cytofix solution, Becton and Dickson, Franklin Lakes, NJ) was added to the cells. Fixed cells were washed 3 times with 200 μl of PBS as before. After washing, the PBS was removed, and cover slips were counterstained and mounted onto microscopic slides using 20 μl DAPI mounting media (VECTASHIELD, Vector Laboratories, Inc. Burlingame, CA). Samples were evaluated by fluorescent microscopy, using Nikon eclipse 80i microscope and the imaging software NIS Elements (Nikon USA, Melville, NY). For confocal experiments samples were prepared as before and evaluated by confocal microscopy using a Zeiss 880 confocal microscope and Fiji software (ImageJ, https://fiji.sc).

### Anti-Cryptosporidial activity of ssRNA/ago

To test for anti-Cryptosporidial activity of ssRNA/Ago *in vitro we quantified parasite reduction on infected cells treated with complexes.* For these experiments we infected cultured HCT-8 cells with *Cryptosporidium* oocysts in 24 well plates as described before (8). Briefly: after excystation, sporozoites were added to HCT-8 cells for 2 hrs to allow infection, then culture media was removed and 250 μl of fresh complete media (RPMI supplemented with 10% FBS and antibiotic) was added and infected cells were incubated at 37°C for 24 hrs. Silencing complexes were assembled and encapsulated in PTR as before and diluted in 100 μl of RPMI media (free serum). Diluted samples were added daily (every 24 hrs) to cultured cells at day 1, 2 and 3 days. Treated cells were incubated and harvested at 24, 48 and 72 hrs, then RNA was extracted with the RNeasy Plus kit following vendor instructions, (Qiagen. Valencia, California). After RNA extraction, samples were transferred to 1.5 ml tubes and stored frozen (−20°C) for subsequent RNA extraction. To evaluate anti-Cryptosporidial of ssRNA/Ago effect on treated cells, total number of parasites was calculated (by detecting 18s Cryptosporidium gene) by RT-PCR in untreated samples and then CT values were used to calculate fold change of treated samples (respect to base line) using the Delta-Delta Ct Method (2^−ΔΔCt^). For RT-PCR assays, CT values were normalized using Gapdh as reference gene. Experiments were conducted in triplicate and data presented as percentage change from base line.

### Labeled complexes delivered to mice intestines

All animal experiments were conducted under a protocol approved by the Institutional Animal Care and Use Committee of the University of Texas Medical Branch. To demonstrate delivery of complexes measured ssRNA uptake on intestinal cells. For these experiments, we as ambled complexes using FAM-labeled ssRNA as and then treated 4-week-old mice with a single dose of PTR containing FAM-labeled ssRNA/Ago (ssRNA [100nM]/Ago [125 ng] diluted in 100 μl of PBS) was given to severe combined immunodeficiency/beige (SCIDbg) mice (Jackson lab, Sacramento, California USA) by oral gavage. PBS or naked FAM-labeled ssRNA/Ago encapsulated in PTR were given by gavage as controls. Mice were sacrificed 4 hours after the oral dose and the terminal ileum was collected. Two to 4 mm sections of ileum were transferred to PBS-EDTA solution and Intestinal crypts were isolated using 2 mM EDTA in PBS as described (19). Crypts were washed twice with PBS and fixed with 4% paraformaldehyde in PBS for 30 min. We quantified cellular uptake of FAM labeled ssRNA/Ago by fluorometric assays as described (20). Briefly: ~ 500 crypts were placed in 5 ml of PBS. Cells were washed twice with PBS and lysed in 400 μl lysis buffer (1% Triton X-100, 2% SDS in PBS) on ice for 30 min. Lysates were centrifuged (15 min, 14,000 g, 4 °C) to remove cell debris. Then, 200 μl of the supernatant was transferred to a black 96-well plate to measure the fluorescence using a BMG FLUOstar Omega Microplate Reader (BMG Labtech Inc. Cary, NC). From sample, 50 μl was used to determine the protein content using the Micro BCA™ protein assay (Pierce, Rockford, USA), according to the protocol of the manufacturer. For determination of the mean fluorescence intensity, fluorescent signals were corrected for the amount of protein in the samples.

### Treatment of infected mice model *in vivo* and parasite quantification

Five-week-old SCIDbg mice (Jackson Laboratory, Sacramento, California) were infected by gavage with 5 × 10^5^ *C. parvum* oocysts (Iowa strain, Waterborne Inc. New Orleans, LA) contained in 100 μL of PBS as described before (21). Four days after infection, mice were treated with ssRNA Ago complexes (prepared as before and suspended in 100 μL of H_2_O) by oral gavage daily for 3 days. Five animals per group were treated with ssRNA-NDK/Ago, ssRNA scramble/Ago or PBS. For parasite quantification, approximately 20 mg of stool (2 pellets) was collected daily and re-suspended in 1 ml of water. The samples were stored at −20°C until subsequent analysis. DNA was extracted and purified from 20 mg (2 pellets) of stool from each mouse using the QIAamp Fast DNA Stool Mini Kit (Qiagen, Germantown, MD). The concentration of DNA in samples was determined by spectrophotometry. The parasite burden was determined by qPCR (Applied Biosystems 7500 Real-Time PCR System), using the iTaq Universal SYBR Green Supermix Kit (Bio-rad, Hercules, California) with 18s primers for *C. parvum* as described (22). The qPCR assay was conducted under the following conditions: 1 cycle of 20 minutes at 55°C, 1 cycle of 5 minutes at 95°C and 15 seconds at 95°C, 40 cycles of 1 minute at 60°C. An additional dissociation stage was added at the end of the reaction to test the specificity via dissociation curve analysis. A standard curve was generated from serial dilutions of DNA extracted from a known number of parasites spiked in mouse stool. Total numbers of parasites were calculated with AB7500 software SDS v1.4.

### Statistical analysis

For all RT-PCR experiments Sigma plot V12 was used for the statistical analysis. Data were analyzed using unpaired two-tailed t test and presented as means ± SD. P < 0.05 was considered to be statistically significant.

## Results

### NDK ssRNA/Ago complexes induce silencing in *Cryptosporidium*

We induced silencing of NDK by transfecting NDK antisense ssRNA/ago complexes into *C. parvum* oocysts (Supplementary, figure S1). Our results showed that complexes targeting NDK mRNA reduced expression by 85% (Fig 1). In contrast naked NDK ssRNA only reduced NDK expression by 65% and sense NDK ssRNA or scramble ssRNA did not show any effect on NDK mRNA levels. Thus, Both, antisense NDK ssRNA, with or without Ago, significantly reduce the expression of NDK. Furthermore, the addition of Ago makes it even more efficient.

**Fig 1.**
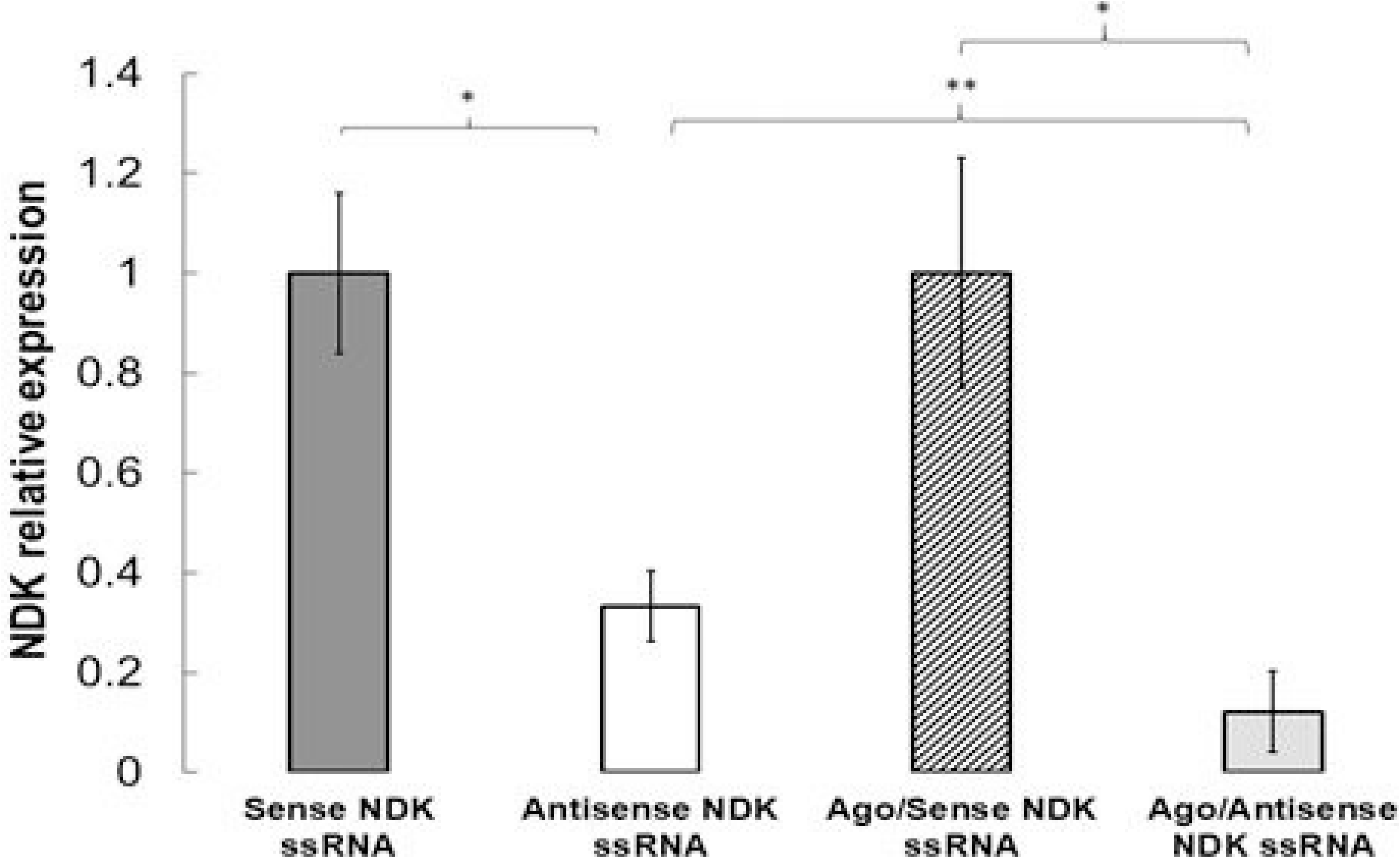
NDK silencing in Cryptosporidium. Oocysts were treated with sense NDK ssRNA (Dark grey), antisense NDK ssRNA (White), Sense NDK ssRNA/Ago (diagonal lines) and antisense NDK ssRNA/Ago (Clear grey). Samples were analyzed by by by qRT-PCR and NDK mRNA expression normalized compared to Cryptosporidium GAPDH and NDK expression compared to untreated controls using the ΔΔCt method. (*P ≤ 0.05, P** ≤ 0.001).

**Figure 2.**
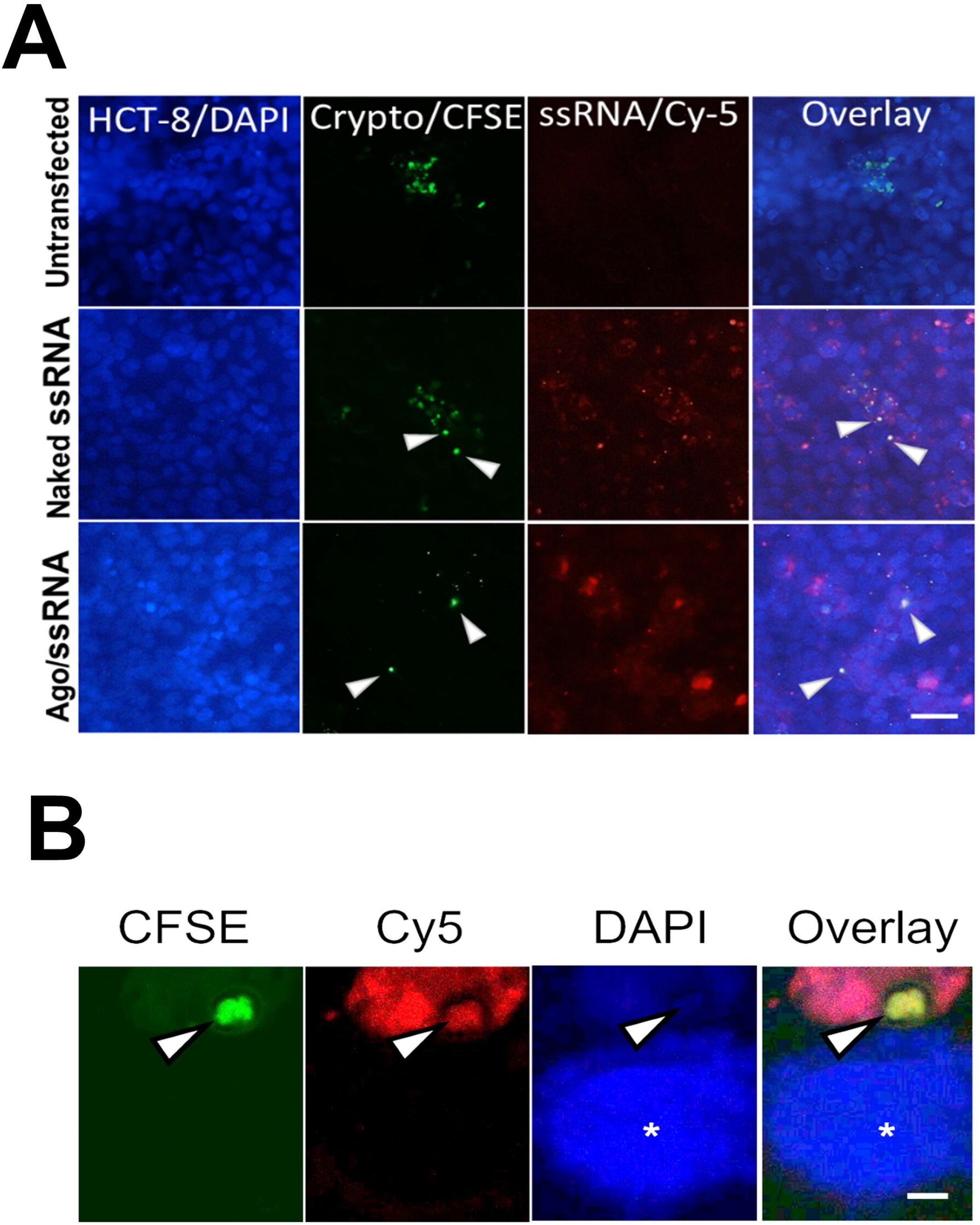
DNK ssRNA/Ago complexes localize to infected cells. HCT-8 cells were infected with fluorescent Cryptosporidium pre-stained with the vital dye CFSE (Green). After 24 hrs. after infection, cells were transfected with Cy-5 labeled ssRNA (Red)/Ago complexes. For fluorescent microscopy (2A), HCT-8 cells were counterstained with DAPI (Blue). White arrows show localization of intracellular parasites. Co-localization is demonstrated ssRNA-Cy5.5 (Naked) and Ago/ssRNA-Cy5.5 (complexes). Scale bar=40 μm. For Confocal microscopy (2B), HCT-8 cells cultured on coverslips were treated as before. Nuclei is stained in blude with DAPI (white *). The white arrow points to an intracellular parasite, which was transfected with ssRNA/Ago complexes. Scale bar =5 μm.

### ssRNA/Ago is delivered to infected cells

We used CFSE-labeled parasites to infect HCT-8 cells. After the infection, we treated intestinal cells with Cy5-ssRNA/Ago encapsulated in liposomes, fluorescent microscopy showed that Cy5-ssRNA/Ago complexes are widely distributed among HCT-8 cells and some of these Cy5 labeled complexes are co-localized with infected cells (2A), also confocal studies confirmed transfection of intracellular parasites (2B). These results suggest that parasitophorous vacuole of infected cells is susceptible to transfection therefore would not be a barrier for treating intracellular parasites with ssRNA/Ago complexes.

### ssRNA/Ago treatment reduces Cryptosporidium infection in vitro

HCT-8 cells were infected with sporozoites. After 24 h, silencing complexes were added daily to infected cells for up to for 3 days. Treatment with silencing complexes quickly reduced the number of parasites as measured by qPCR. There was a 75% reduction in parasite mRNA after day 1 and 95% by 3 days of treatment (Fig 3). Cells treated with naked RNA only had a 50% reduction in parasite numbers by day 3 (Fig 3). These results confirm that Ago enhances the potency of silencing and suggest the feasibility of using complexes as potential treatment of cryptosporidiosis in vivo.

**Figure 3.**
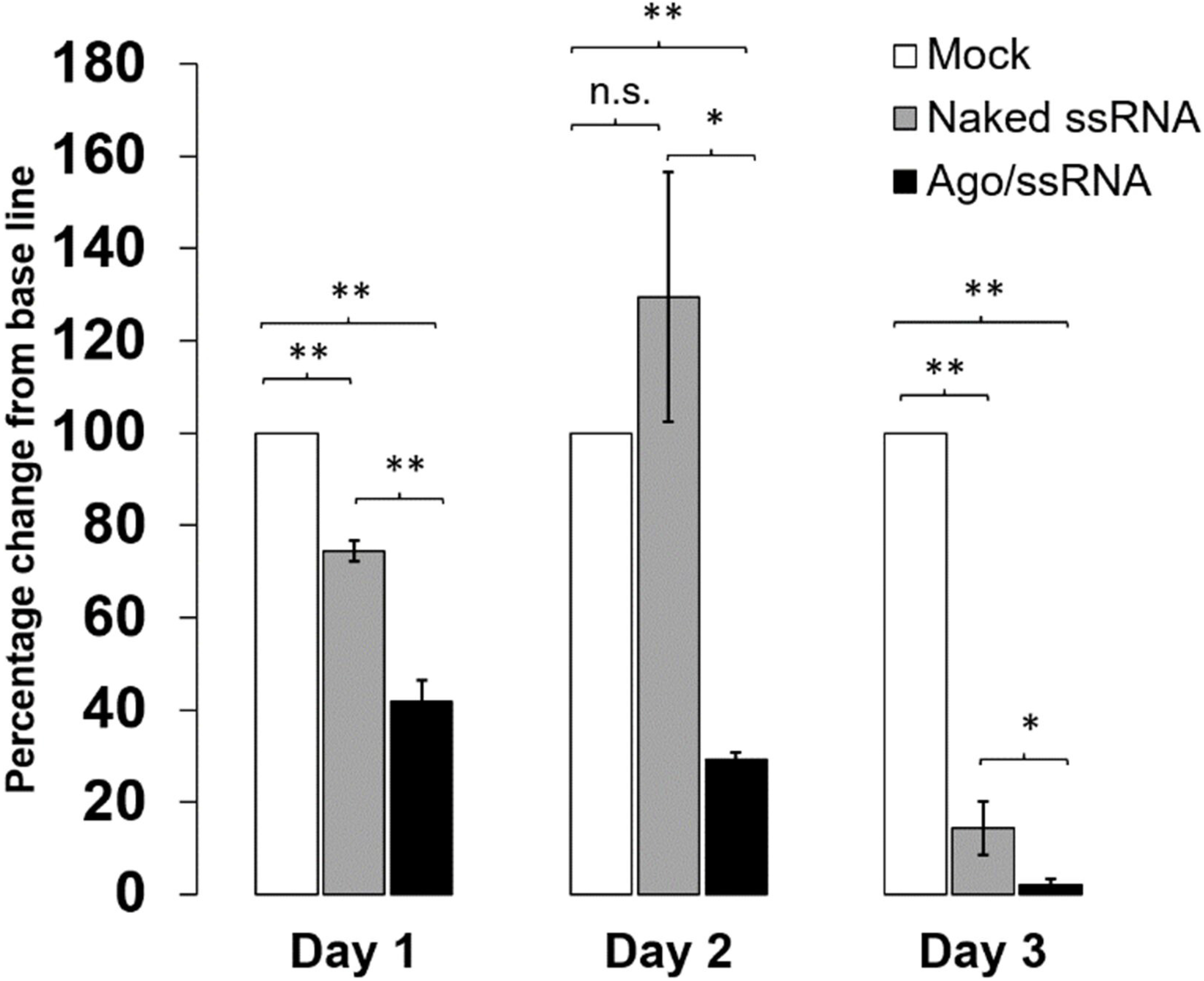
In vitro Anti-Cryptosporidial activity of ssRNA/Ago. HCT-8 cells were infected Cryptosporidium parvum. After 24 hrs, infected cells were treated daily with anti NDK ssRNA (grey bar) or anti-NDK ssRNA/Ago complexes (black bars). Control infected cells were treated with scramble ssRNA (Mock). Total number of parasites (base line=100%) was calculated by qRT-PCR and results in triplicate are expressed as percentage change from base line. (*P ≤ 0.05, P** ≤ 0.005).

### ssRNA complexes are delivered to mouse intestine

We investigated whether ssRNA/Ago complexes can reach the mouse intestines after oral administration. For these experiments we treated SCIDbg mice with labeled ssRNA and then we analyzed ssRNA/Ago uptake with or without lipidic encapsulation on intestinal cells by fluorometry (Fig 4). Both, naked ssRNA/ago and complexes were detected on mice intestinal cells 4 hrs. after treatment. However, Uptake experiments showed a stronger signal with encapsulated ssRNA/Ago (Fig 4). These results show the feasibility to use of lipidic complexes to enhance delivery of ssRNA/Ago on mice intestines.

**Figure 4.**
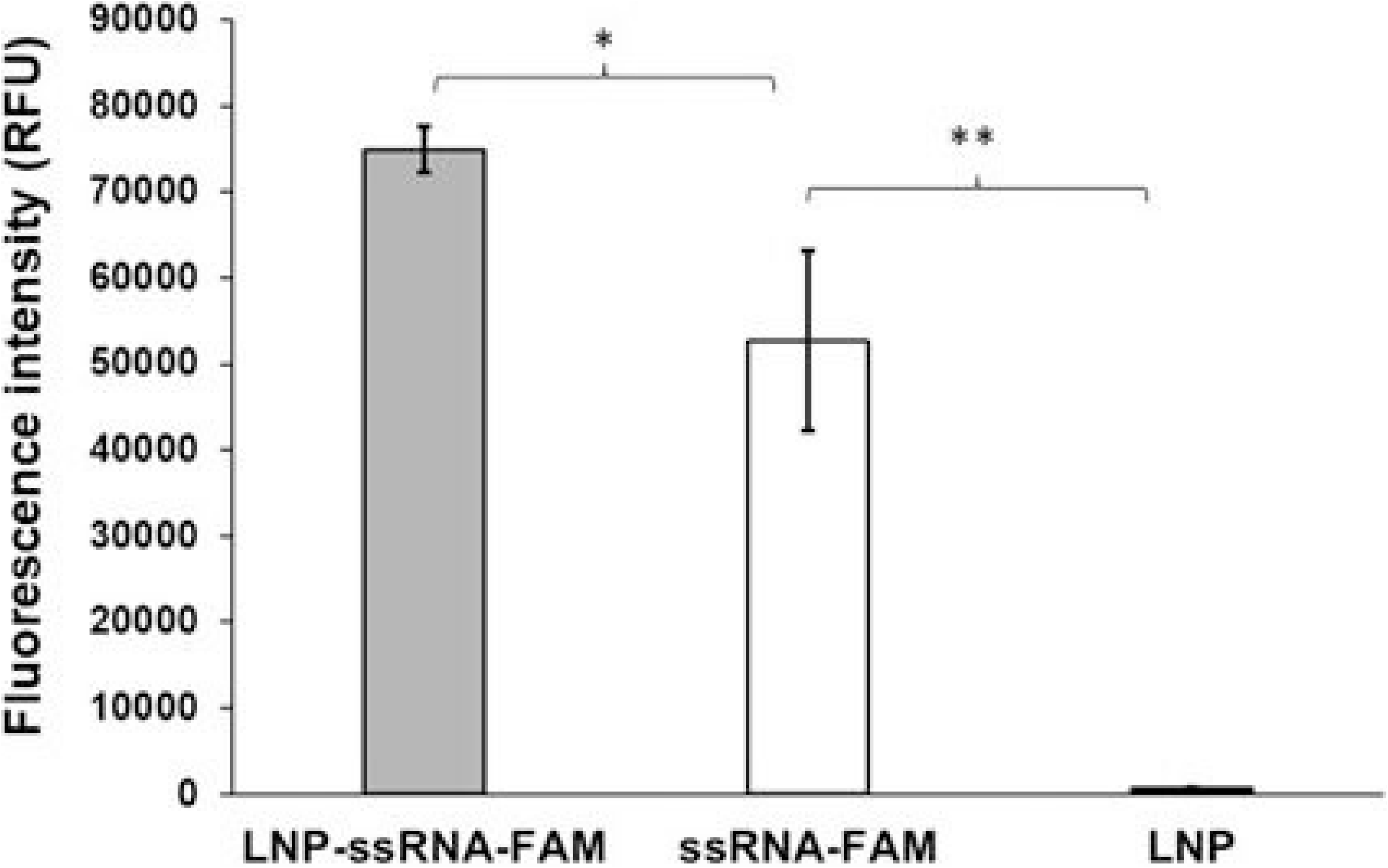
Intestinal cells of mice treated with labeled ssRNA. Mice treated orally with labeled ssRNA-FAM/Ago complexes encapsulated in lipid-based nanoparticles (LNP), after 4 hrs intestinal cells were obtained and cellular uptake analyzed by fluorometry. Fluorescence intensity is measured in relative fluorescence units (RFU). Grey bar: mice treated with ssRNA/Ago complexes encapsulated in LNP, white bar: mice treated with ssRNA/Ago complexes without LNP, Black bar: mice treated only with LNP. (*P ≤ 0.05, P** ≤ 0.001).

### Anti NDK ssRNA reduces the parasite burden in infected mice

We evaluated the anti-cryptosporidal activity of ssRNA/Ago complexes in SCIDbg mice infected with *Cryptosporidium*. After confirming oocyst shedding, mice were treated by gavage with anti NDK/Ago or scramble NDK/ago beginning on day 4 post infection. Four of 5 mice treated with NDK/Ago cleared the infection by day 3 (Fig 5). In contrast, 4/5 mice treated with scrambled ssRNA/Ago continued to shed oocysts at levels similar to untreated mice (Fig. 5). This experiment confirms that NDK ssRNA/ago has anti-cryptosporidial activity and suggest the feasibility of using antisense therapy to treat cryptosporidiosis.

**Figure 5.**
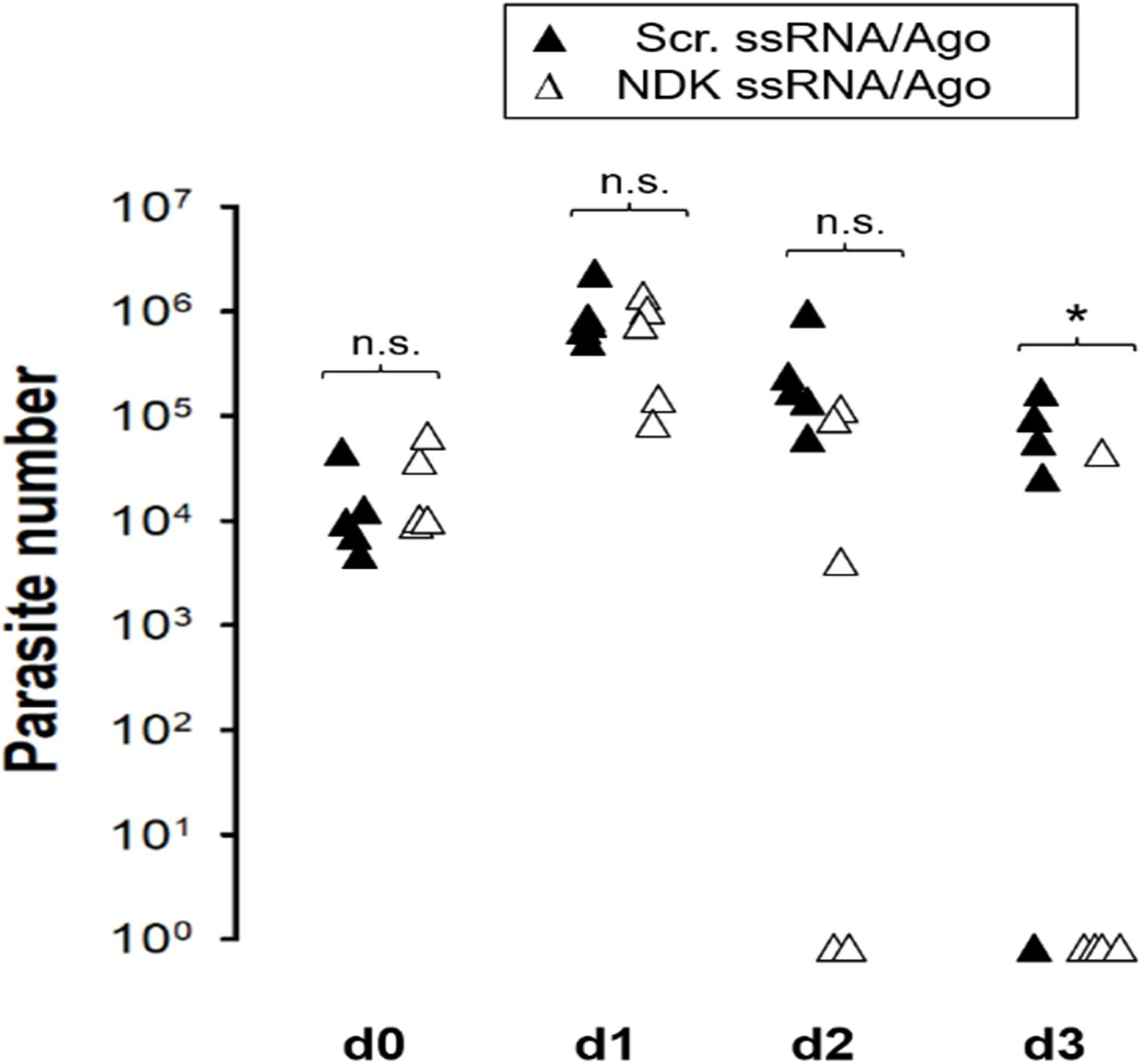
Parasite burden in stool samples of mice orally treated with ssRNA/Ago. Mice were infected with 1×106 parasites, after 4 days of infection mice were treated orally with daily with ssRNA/Ago complexes encapsulated in LPN. Parasite burden in 25 mg of stool was evaluated by qPCR. Anti-NDK ssRNA/Ago complexes (black triangles), scramble ssRNA/Ago (white triangles). *=P ≤ 0.05.

## Discussion

We have induced gene silencing in *Cryptosporidium* using preformed “ssRNA/Ago complexes and showed that knocking down NDK expression blocks parasite proliferation (8). Here, we demonstrated the feasibility for delivering ssRNA/Ago complexes to the intestines of mice and demonstrated anti-Cryptosporidial activity of antisense NDK ssRNA/Ago complex in vitro and *in vivo*. *Cryptosporidium* lacks RISC enzymes (23), therefore our silencing strategy is based in the reconstitution of the small interfering RNA pathway (siRNA) by forming a minimal RNA silencing complex (7).

Recent studies have shown that siRNAs, can be used therapeutically to block the synthesis of disease-causing proteins and also to reduce proliferation of infectious agents (24). However, the delivery of nucleic acids by the oral route poses potential hurdles, including instability of the RNA or poor cellular uptake could compromise efficacy under physiological conditions. In addition, high concentrations of siRNA could also induce off-target effects or nonspecific inflammatory responses (25). We hypothesized that preformed ssRNA/Ago complexes could induce silencing at lower concentrations. In vitro studies confirmed that low concentrations of ssRNA preloaded onto Ago induce potent silencing in transfected parasites. Previous studies have shown that Ago binding protects ssRNA from degradation (26). We demonstrated that oocysts (supplementary figure 1S) and intracellular parasites are susceptible to transfection using a cationic lipid-based carrier system, is known that this kind of liposomes attach to negatively charged surfaces and fuse directly to membranes. Therefore, since *Cryptosporidium* do not have caveolins, we speculate that ssRNA/Ago complexes are introduced by Clathrin-mediated endocytosis where the liposomes are fused with parasitophourus membrane and membranes of extracellular forms. Our confocal studies confirmed that ssRNA/Ago complexes cross membranes of intracellular and extracellular parasites (Fig 2B and S1) however additional studies should be conducted to understand how ssRNA/Ago complexes cross membranes on infected cells. For silencing experiments, we targeted Cryptosporidium nucleoside diphosphate kinase (NDK), our previous studies have shown that NDK silencing with ssRNA/Ago complexes blocks parasite proliferation without cytotoxic effects. Our in vitro experiments Our confirmed these observations and showed that a daily treatment (up to 3 days) enhances anti-Cryptosporidial activity. Since previous studies have shown that siRNA encapsulated in lipidic nanoparticles is efficiently delivered orally on mice intestines, then we hypothesized that ssRNA/Ago could be delivered in vivo using same cationic lipids used in our in vitro experiments. Uptake experiments showed on fig. 4 confirmed that cationic lipids enhance delivery of complexes on epithelial cells, suggesting the feasibility to use NDK ssRNA/Ago to reduce infection in Vivo. Since this concept has not had tested before, then we conducted a preliminary experiment using a *Cryptosporidiosis*-mouse model to confirm if daily treatment with NDK ssRNA/Ago (using same doses) showed anti-Cryptosporidial activity on infected mice. Our results showed that complexes were detected in the mouse intestinal tissues and led to significant reduction of parasites by day 2 in all mice with 4 of 5 mice showing no infection by day 3 (Fig. 5). However, further optimization of dose, schedule, and duration of therapy may be needed prior to initiating studies in large animal models. NDK is primarily involved in nucleotide synthesis. However, recent genetic experiments in this parasite showed that under certain conditions *Cryptosporidium* may not require purine nucleotide synthesis (27). Those studies proposed that the parasite has evolved to import purine nucleotides, making purine salvage dispensable. Thus, the effect of NDK inhibition might be related with other NDK functions. NDK has also been implicated in the evasion of immune response, apoptosis and inflammation.

*Proprionobacterium gingivalis*-NDK inhibits extracellular-ATP (eATP)/P2X7-receptor mediated cell death in gingival epithelial cells (GECs) via eATP hydrolysis (28). For *L. amazonensis*, NDK participates in immune evasion by preventing eATP-mediated macrophage death (13). Specifically, *Leishmania*-secreted NDK was found to keep the host cell membrane integrity intact, and stabilized the mitochondrial membrane potential in macrophages, thus preventing host-cell death. (12).

Other groups have been actively involved in drug discovery efforts for cryptosporidiosis (3, 29). Phenotypic screening of repurposed drugs has identified several potential lead compounds (30). However, one important lead, clofazimine, was effective *in vitro* and in small animal models, but had no activity in clinical trials (31). Other research groups have worked on development of novel drugs for parasite targets. Significant efforts have targeted the parasite calcium dependent protein kinase 1 (32). A number of highly effective compounds have been developed. However, so far, all have failed to progress to clinical trials due to unanticipated adverse effects. Additional avenues are needed for novel drugs for cryptosporidiosis. Our work is based in RNA interference technology. Compared to other small molecule drugs or antibody-based treatments, siRNA can potentially have improved specificity determined by complementary base pairing. Recent studies have demonstrated the feasibility to use RNAi against infectious agents by using different strategies to enhance potency, stability, and delivery of nucleic acids (24). Our preliminary studies test for first time a novel strategy based on the use of argonaute and protein transfection reagents to conduct in vivo silencing. Overall, this work shows the feasibility to use ssRNA/Ago complexes for treatment of cryptosporidiosis.

